# MinION-based DNA barcoding of preserved and non-invasively collected wildlife samples

**DOI:** 10.1101/2020.01.29.925081

**Authors:** Adeline Seah, Marisa C.W. Lim, Denise McAloose, Stefan Prost, Tracie Seimon

## Abstract

1. The ability to sequence a variety of wildlife samples with portable, field-friendly equipment will have significant impacts on wildlife conservation and health applications. However, the only currently available field-friendly DNA sequencer, the MinION by Oxford Nanopore Technologies, has a high error rate compared to standard laboratory-based sequencing platforms and has not been systematically validated for DNA barcoding accuracy for preserved and non-invasively collected tissue samples.
2. We tested whether various wildlife sample types, field-friendly methods, and our clustering-based bioinformatics pipeline, SAIGA, can be used to generate consistent and accurate consensus sequences for species identification. Here, we systematically evaluate variation in cytochrome b sequences amplified from scat, hair, feather, fresh frozen liver, and formalin-fixed paraffin-embedded (FFPE) liver. Each sample was processed by three DNA extraction protocols.
3. For all sample types tested, the MinION consensus sequences matched the Sanger references with 99.29-100% sequence similarity, even for samples that were difficult to amplify, such as scat and FFPE tissue extracted with Chelex resin. Sequencing errors occurred primarily in homopolymer regions, as identified in previous MinION studies.
4. We demonstrate that it is possible to generate accurate DNA barcode sequences from preserved and non-invasively collected wildlife samples using portable MinION sequencing, creating more opportunities to apply portable sequencing technology for species identification.

## Introduction

Wildlife health and conservation initiatives benefit tremendously from genetic methods of species identification for infectious disease screening (Schlaberg, Chiu, Miller, Procop, & Weinstock, 2017; Gardy & Loman, 2018), detecting illegally traded wildlife products (Hobbs, Potts, Walsh, Usher, & Griffiths, 2019), uncovering food label fraud (Pardo et al., 2018; Galimberti et al., 2019; Hobbs et al., 2019), and documenting understudied biodiversity (Costa & Carvalho, 2007). One major challenge for wildlife molecular studies is obtaining fresh samples from live or dead wild animals. Such endeavors can be logistically challenging, generally involving highly skilled teams, detailed planning, and acquisition of permissions from local, regional and international partners and governmental agencies for animal handling, sample collection, and sample transfer for molecular testing. Consequently, environmental samples (Ficetola, Miaud, Pompanon, & Taberlet, 2008; Thomas et al., 2019) and animal samples that can be collected non-invasively (e.g. hair, feathers, scat, etc.) (Marshall & Ritland, 2002; Waits & Paetkau, 2005; De Barba et al., 2014) are increasingly being used for ecological studies, wildlife health assessments, and characterizing biodiversity. Non-invasively collected samples are easier to obtain than fresh organ tissues, but may contain PCR inhibitors, have lower DNA yields, or are degraded from environmental exposure (Kohn, Knauer, Stoffella, Schröder, & Pääbo, 1995; Rådström, Knutsson, Wolffs, Lövenklev, & Löfström, 2004; Waits & Paetkau, 2005; Chaturvedi et al., 2008). Archived historical wildlife samples, often preserved in formalin, also offer a unique opportunity to obtain genetic information (Seimon et al., 2015). However, challenges for molecular studies include formalin-related fragmentation and DNA cross-linking (Do & Dobrovic, 2015; Einaga et al., 2017).

DNA barcoding is a common molecular technique for species identification (Hebert, Ratnasingham, & de Waard, 2003; Valentini, Pompanon, & Taberlet, 2009). The Oxford Nanopore Technologies (ONT) MinION sequencer is currently the only available portable sequencer. Although nanopore sequencing is known to have higher raw sequence error rates in comparison to standard short read sequencing platforms such as Illumina or BGI-Seq, particularly at homopolymeric regions (Ip et al., 2015; Jain et al., 2017), significant improvements in the accuracy of MinION sequencing chemistry has led to its recent rise in popularity for field applications (reviewed in Krehenwinkel, Pomerantz, & Prost, 2019). This sequencer is especially useful in situations where there is a lack of access to sequencing facilities or when sample export is difficult. The MinION also has a lower investment cost and shorter turnaround times than traditional sequencing platforms (e.g., Sanger, Illumina).

MinION DNA barcoding studies have primarily used laboratory-based QIAGEN® kits for reliable and pure DNA extraction products (e.g., Pomerantz et al., 2018; Krehenwinkel, Pomerantz, Henderson, et al., 2019; Maestri et al., 2019). To expand the potential for portable sequencing applications, field-friendly DNA extraction methods can be used to reduce lab equipment requirements. While field-friendly DNA extraction methods are often less effective at producing DNA of high concentration and purity levels, MinION DNA barcoding has been successfully performed using QuickExtract™ solution (Lucigen), which only requires a heat source (Srivathsan et al., 2019). The Chelex® 100 resin (Bio-Rad Inc.) extraction method similarly only requires a heat source, but is less expensive and has not been tested for MinION sequencing so far. Both methods have short protocols, but do not remove cellular debris or PCR inhibitors, which can affect downstream applications (Walsh, Metzger, & Higuchi, 1991; Singh, Kumari, & Iyengar, 2018). The Biomeme M1 Sample Prep™ Kit (Biomeme Inc.) is another DNA extraction kit developed for field use. While more expensive than either QuickExtract or Chelex methods, the Biomeme kit includes all necessary components and both protein and salt wash steps to remove impurities. Studies have shown that Biomeme-extracted samples have higher levels of inhibitors compared to Qiagen extractions, and thus requires additional dilution steps (Sepulveda, Hutchins, Massengill, & Dunker, 2018; Thomas et al., 2019).

To date, MinION DNA barcoding pipelines have used either *de novo* assembly (Pomerantz et al., 2018; Krehenwinkel, Pomerantz, Henderson, et al., 2019), clustering-based (Maestri et al., 2019), or alignment (Srivathsan et al., 2018, 2019) methods to generate consensus sequences for species identification. Assembly approaches generally work more consistently for longer barcodes (~1kb), as the underlying software were originally designed for assembling long reads for genome assemblies rather than amplicons. Both published clustering or alignment pipelines use subsets of the data (100-200 reads) to generate scaffolds for read error correction. While these approaches may work for high quality sequence data, the data subsets could include more sequence error bias in lower quality datasets. Thus, we developed a clustering-based pipeline, SAIGA (https://github.com/marisalim/Saiga), with software specifically designed for error prone MinION reads that processes data regardless of barcode length, and maximizes the use of demultiplexed reads for downstream species identification analysis.

In this study, we systematically evaluate the accuracy of the MinION for DNA barcoding across a range of wildlife sample types, including two field-friendly DNA extraction approaches. We sequenced a short fragment of the commonly used mitochondrial cytochrome b (Cytb) gene from scat, hair, feather, fresh frozen liver and formalin-fixed paraffin embedded (FFPE) liver. For each sample type, we compared the accuracy of Cytb consensus sequences for three different DNA extraction methods: QIAGEN silica membrane-based kits, Chelex 100 resin, and the Biomeme M1 Sample Prep Kit. All analyses were conducted with SAIGA. We demonstrate that MinION sequencing can be used with field-friendly extraction methods to accurately identify wildlife species from a variety of sample types.

## Materials and Methods

### Sample collection

For this study, scat, hair, feather, fresh frozen liver and FFPE liver samples were collected opportunistically during necropsy examinations from a snow leopard (*Panthera uncia*) and a cinnamon teal (*Anas cyanoptera*) from a zoological collection. The FFPE liver samples were part of a suite of tissues that were collected, stored in 10% neutral buffered formalin, and subsequently processed and paraffin-embedded for histologic examination and routine tissue archiving. Fresh liver, scat, hair and feather samples were frozen (−80°C) immediately after collection.

### DNA extraction

DNA was extracted from each sample type using three different approaches: 1) Qiagen (QIAamp® DNA minikit or QIAamp® DNA Stool Mini Kit, Qiagen Inc., Germantown, MD, USA); 2) Chelex 100 Resin (Bio-Rad, Hercules, CA, USA); and 3) Biomeme M1 Sample Prep Kit for DNA (Biomeme, Philadelphia, PA, USA). DNA quantification is inaccurate for Chelex extracts due to the presence of cellular components, thus Chelex extracts were not quantified. All Qiagen and Biomeme extracts were quantified using the Qubit™ dsDNA High Sensitivity Kit on the Qubit™ 4 Fluorometer (Thermo Fisher Scientific, Waltham, MA, USA). The Qiagen, Chelex, and Biomeme extraction protocols are summarized for each tissue type in Appendix I. All Qiagen, Biomeme DNA extracts with >10ng/μL, and all Chelex extracts were run on a 1.0% gel to assess DNA fragmentation by sample type.

### PCR & library preparation

#### DNA Barcoding PCR - Round 1

Approximately 460 bp of the mitochondrial Cytb gene was amplified using primers mcb398 and mcb869 (Verma & Singh, 2003), with universal tailed sequences on each primer that are compatible with the ONT PCR Barcoding Expansion kit EXP-PBC001 (ONT, Oxford, UK) (Table S1). These primers were designed from an alignment of 67 animal species, and validated for mammals, reptiles and birds (Verma & Singh, 2003).

PCR was carried out with 6.25 μL DreamTaq HotStart PCR Master Mix (Thermo Fisher, Waltham, MA, USA), 1.25 μL DNA template, and 2 μL of each primer (10 μM stock) in a final volume of 12.5 μL. Cycling conditions were: 95°C for 3 minutes; 35 cycles of 95°C for 30 seconds, 55°C for 30 seconds and 72°C for 30 seconds; and a final extension of 72°C for 5 minutes. All Chelex extractions were diluted for the DNA Barcoding PCR as described in Appendix I. PCR products were purified using 1.8X Agencourt AMPure XP beads (Beckman Coulter, Indianapolis, IN, USA), tested for purity using the NanoDrop™ One spectrophotometer (Thermo Fisher Scientific, Waltham, MA, USA), and quantified fluorometrically using the Qubit dsDNA High sensitivity kit.

#### Indexing PCR - Round 2

To attach dual ONT PCR index sequences to the Cytb amplicons, a second round of PCR was carried out with the ONT PCR Barcoding Expansion kit for each sample with 25 μL KAPA Biosystems HiFi HotStart ReadyMix (2X) (Thermo Fisher Scientific, Waltham, USA), containing 25 ng of first-round PCR amplicon and 1 μL ONT PCR Barcode in a final volume of 50 μL. Cycling conditions were: 95°C for 3 minutes; 11 cycles of 95°C for 15 seconds, 62°C for 15 seconds and 72°C for 15 seconds; and a final extension of 72°C for 1 minute. Hereafter, we refer to ONT PCR barcodes as ‘indexes’ to reduce confusion with the Cytb barcode. Indexed PCR products from round 2 were purified and tested for purity and quantity like round 1 products.

#### Library preparation

Samples were grouped into four libraries by sample type (FFPE, scat, hair/feather, frozen liver). For each library, purified indexed amplicons were pooled in equal ratios to produce 1.0-1.2 μg in a total of 45 μL nuclease-free water. Pooled libraries were next prepared using the ONT Ligation Sequencing kit SQK-LSK109 (ONT, Oxford, UK) with modifications to the manufacturer’s instructions: 25 μL of the pooled library was mixed with 3.5 μL NEBNext Ultra II End-Prep Reaction buffer and 1.5 μL Ultra II End-prep Enzyme mix (New England Biolabs, Ipswich, MA, USA), incubated for 10 minutes at room temperature, then 10 minutes at 65°C. For adapter ligation, 15 μL of the end-prepped library (not bead-purified) was mixed with 25 μL Blunt/TA Ligase and 10 μL Adapter Mix (AMX), incubated at room temperature for 20 minutes and eluted in a final volume of 12 μL of Elution Buffer.

### Sequencing

The four libraries were split between two FLO-MIN106D R9.4.1 chemistry flow cells (ONT, Oxford, UK) - to minimize bleed-through between experiments - FAL19910: 1) FFPE, 2) scat; FAL19272: 1) hair/feather, 2) frozen liver. Flow cells were washed with Wash Solution A followed by the addition of Storage buffer S according to the manufacturer’s protocols. All libraries were sequenced for approximately 1 hour to obtain at least 100,000 raw reads per sample.

For comparison to MinION sequences, Sanger sequencing in the forward and reverse directions was performed on all purified indexed amplicons (Eton Bioscience Inc. Newark, NJ, USA). Sanger consensus sequences were generated using Geneious Prime v2019.0.4 software (Biomatters LDT, Auckland, NZ).

### Bioinformatics

The SAIGA bioinformatics pipeline is available on GitHub (https://github.com/marisalim/Saiga) and steps are outlined in Fig. 1. MinKNOW (ONT) was used for sequencing and the raw sequence data were basecalled using Guppy v3.5.1 (ONT) with basecalling model “dna_r9.4.1_450bps_fast.cfg”.

**Figure 1:**
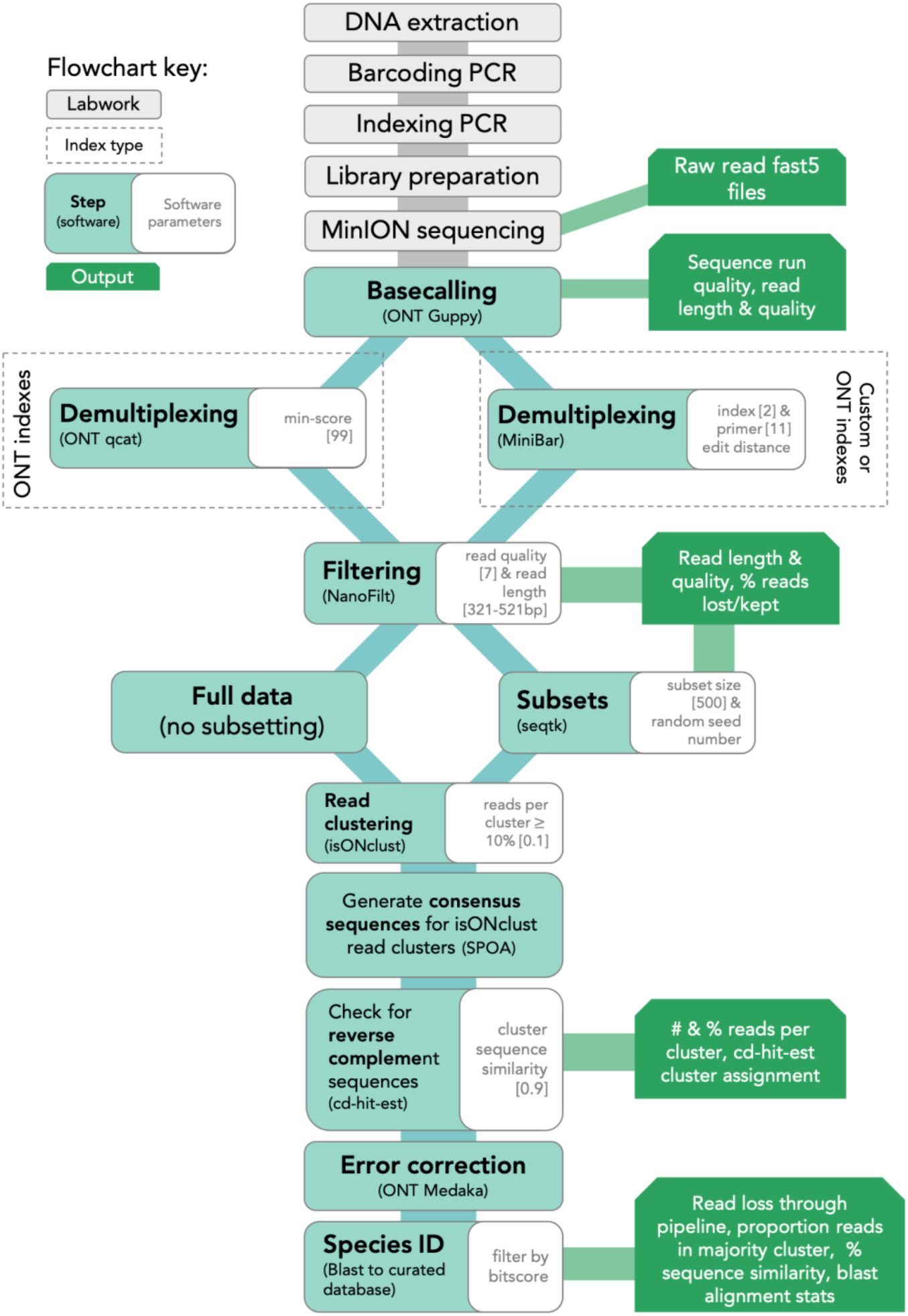
Lab and SAIGA bioinformatics pipeline flowchart. Bioinformatics software and parameters are indicated at each step.

#### Demultiplexing and filtering

Assigning sequencing reads to the correct sample is a critical step to avoid mixing sample sequences within or between sequencing runs. Thus, we compared results from two demultiplexing programs: 1) qcat v1.1.0 (ONT, https://github.com/nanoporetech/qcat) and 2) MiniBar v0.21 (Krehenwinkel, Pomerantz, Henderson, et al., 2019). The qcat software was built specifically for demultiplexing reads indexed with ONT’s barcode kits, while MiniBar is a general demultiplexing software that allows any set of user-specified index and primer sequences. We used stringent demultiplexing filters based on software recommendations, sensitivity analyses, and to minimize incorrect read assignments. Qcat uses the epi2me demultiplexing algorithm and we trimmed adapter and index sequences with the trim option. Using the min-score option, demultiplexed reads with alignment scores <99 were removed prior to downstream analysis, where a score of 100 means every nucleotide of the index is correct. Lower min-score thresholds (i.e., 60-90) reduced downstream consensus sequence quality. In MiniBar, up to 2 nucleotide differences between reads were allowed for the index sequences and 11 nucleotide differences between primer sequences per software recommendations; MiniBar primarily uses the index sequence information to demultiplex and trim dual index and primer sequence.

After demultiplexing, reads were removed if they had mean Phred quality scores <7 and were longer or shorter than the target amplicon length (~421 bp excluding primers) with a 100 bp buffer (321-521 bp) in NanoFilt v2.5.0 (De Coster, D’Hert, Schultz, Cruts, & Van Broeckhoven, 2018). Following each of the above steps, we calculated and visualized read quality statistics for raw, demultiplexed, and filtered reads with NanoPlot v1.21.0 (De Coster et al., 2018). To standardize dataset size across the four sequencing experiments and to investigate the effect of read depth, we generated 100, 500, and 5,000 random read subsets for each sample from the filtered demultiplexed read files. Hereafter, we refer to these subsets as 100R, 500R, and 5KR, respectively.

#### Read clustering and consensus sequence generation

To generate the consensus sequence for each sample, all reads were first clustered using isONclust v0.0.4 (Sahlin & Medvedev, 2018). We chose isONclust over clustering tools previously used in nanopore-based DNA barcoding pipelines, such as VSEARCH (implemented in ONTrack, Maestri et al., 2019), as it was specifically designed to work with error-prone long-read data and thus should be less affected by read errors and more efficient in cluster formation. Next, SAIGA outputs the number of reads per cluster, only retaining clusters with >10% of the total reads (user-defined). We implemented this step to minimize the inclusion of reads with high sequence error and possible contaminant reads in downstream analysis. Intermediate consensus sequences are then generated using SPOA v3.0.1 (https://github.com/rvaser/spoa), which is based on a partial order alignment (POA) algorithm (Lee, 2003). SPOA also conducts error corrections, resulting in more accurate consensus sequences. The SPOA consensus sequences are then clustered using cd-hit-est v4.8.1 with a stringent similarity cutoff (0.9; user-defined) (Li & Godzik, 2006; Fu, Niu, Zhu, Wu, & Li, 2012). Since isONclust separates reads in different strand orientations, this second round of clustering groups reverse-complement SPOA consensus sequences, ensuring that more filtered reads are used for generating the final consensus sequence. The reads contributing to all SPOA consensus sequences that group with the majority isONclust cluster’s SPOA consensus sequence are combined into a single file for mapping. SAIGA then maps these reads to the SPOA consensus sequence of the majority isONclust cluster for consensus polishing with ONT’s Medaka software v0.10.0 (https://github.com/nanoporetech/medaka).

### Consensus accuracy and analysis

The MinION consensus sequences were compared to Sanger sequences from the same sample using a nucleotide Blast search v2.8.1+ (Altschul, Gish, Miller, Myers, & Lipman, 1990). To assess and compare species identification results across tissue types, extraction methods, demultiplexing programs, and data subsets, the following were evaluated: 1) the percent of matching nucleotides between consensus and Sanger sequences, 2) the number of matching nucleotides between consensus and Sanger sequences, and 3) the proportion of filtered reads in the cluster used to generate final consensus sequence. Accurate species identification was defined as those with >99% sequence similarity to the Sanger sequence and ~421 bp of matching nucleotides. The proportion of demultiplexed reads contributing to the final consensus indicates how much data was used for species identification. For samples with consensus sequences generated from fewer than ~75% of reads, we investigated the non-majority isONclust clusters for potential sequence error or contaminant reads. Finally, all MinION consensus and Sanger sequences across tissue types, extraction methods, demultiplexing software, and data subsets were aligned with Mafft v1.3.7 in Geneious Prime v2019.0.4 to identify common regions with sequence errors.

## Results

### DNA Barcoding and Indexing PCR performance

DNA concentrations were higher for Qiagen (0.8 to 59 ng/μL, n=8) compared to Biomeme (0.07 to 13.9 ng/μL, n=8) extractions (Table S2); Chelex samples were not quantified (n=8). Gel electrophoresis of Qiagen-extracted tissues show frozen liver and scat samples had high molecular weight genomic DNA, while FFPE samples were fragmented; hair and feather extracts were too faint to detect reliably. (Fig. S1). We were unable to detect high molecular weight nucleic acid in the Biomeme and Chelex-extracted samples (Fig. S2). Despite variation in starting DNA concentration and the presence of low molecular weight fragments in some samples, we successfully barcoded and indexed 22 of 24 samples. The two samples that failed to amplify at the Barcoding PCR (Round 1) step were the snow leopard FFPE samples extracted by the Chelex and Biomeme protocols. The DNA concentration of DNA Barcoding PCR (Round 1) products after bead clean-up was <13.9 ng/μL with an average of 3.49 ng/μL. At these low DNA concentrations, NanoDrop purity of Barcoding Round 1 amplicons is highly variable and not reliable.

Two samples had less than 25 ng for Indexing PCR (Round 2). After bead clean-up, the concentration of the snow leopard liver/Chelex DNA Barcoding PCR (Round 1) product was much lower than expected (4.4 ng), despite having a bright agarose gel band. Nevertheless, this was sufficient for amplification in the Indexing PCR step. Cytb was also difficult to amplify from the snow leopard scat/Chelex, so amplicons from two DNA Barcoding (Round 1) PCR reactions were pooled for a total of 16 ng to proceed with Indexing PCR (Round 2). After the Indexing PCR (Round 2) bead clean-up, DNA concentrations were >19 ng/μL with an average of 80.92 ng/μL for all but the snow leopard liver/Chelex sample, which had 6.58 ng/μL. Average A260/280 ratios (1.82) and A260/230 ratios (1.96) indicated relatively pure samples for library preparation.

### MinION and Sanger sequencing performance

Sequencing efficiency, also called pore occupancy, ranged from 72-80% and was evenly spread across flow cells for all MinION sequencing runs (Fig. S3). We sequenced an average of ~752,856 raw reads per run, with an average read length of ~597 bp and read quality Phred score of 10.5 (Table S3, Fig. S4).

We obtained clean Sanger sequences for 21 of 22 samples, all of which were 421 bp after primer trimming (Table S4). For all 21 samples, the Sanger sequences for each species were identical, regardless of tissue type or extraction method. We were unable to get a clean Sanger sequence for the snow leopard scat/Chelex sample. Therefore, we compared the MinION scat/Chelex consensus to the Sanger sequences from the other snow leopard samples for species identity.

### Sequence read retention after demultiplexing and filtering

The average read quality and read lengths were similar across all samples demultiplexed with MiniBar or qcat (Table S3-S4). For all sequencing runs, both MiniBar and qcat correctly assigned demultiplexed reads only to the ONT indexes used in the Indexing PCR for each run (Fig. 2). Due to the stringent demultiplexing thresholds, the majority of read data loss occurred during the demultiplexing step (84.07% reads lost on average; Table S3). After read quality and length filtering, we retained nearly all demultiplexed reads (95.6% reads retained on average; Fig. S5, Table S3). On average, samples had more than 20,000 demultiplexed and filtered reads for downstream analyses (Table S4). In general, MiniBar-demultiplexed datasets retained more reads than qcat-demultiplexed datasets after filtering (Fig. S5). The only sample that retained fewer than 90% of reads after filtering was the cinnamon teal scat/Biomeme sample demultiplexed with MiniBar (68.90% reads retained).

**Figure 2:**
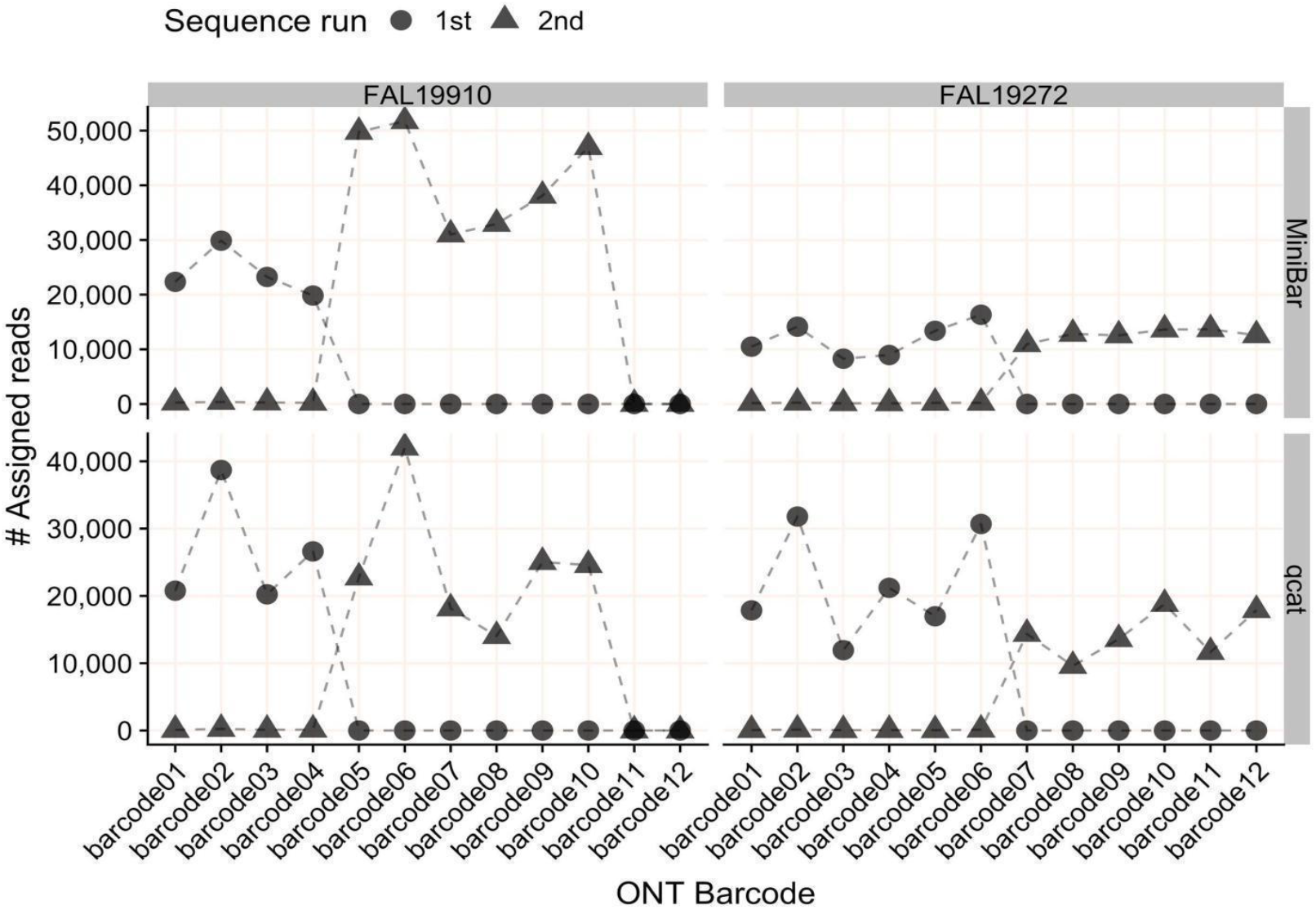
The number of reads assigned to each ONT index (01-12) per flow cell by MiniBar and by qcat. For flow cell FAL19910, the 1st sequencing run used indexes 01-04 and the 2nd run used indexes 05-10. For flow cell FAL19272, the 1st sequence run used indexes 01-06 and the 2nd run used indexes 07-12.

### Read clustering proportions and cluster species identity

For nearly all data subsets, there were only two isONclust clusters for each sample comprising forward and reverse-complement oriented reads. In these cases, 100% of filtered reads formed a single cluster after cd-hit clustering (to merge potential reverse-complements) and all reads were used to produce the consensus sequence for final species identification (Fig. 3).

**Figure 3:**
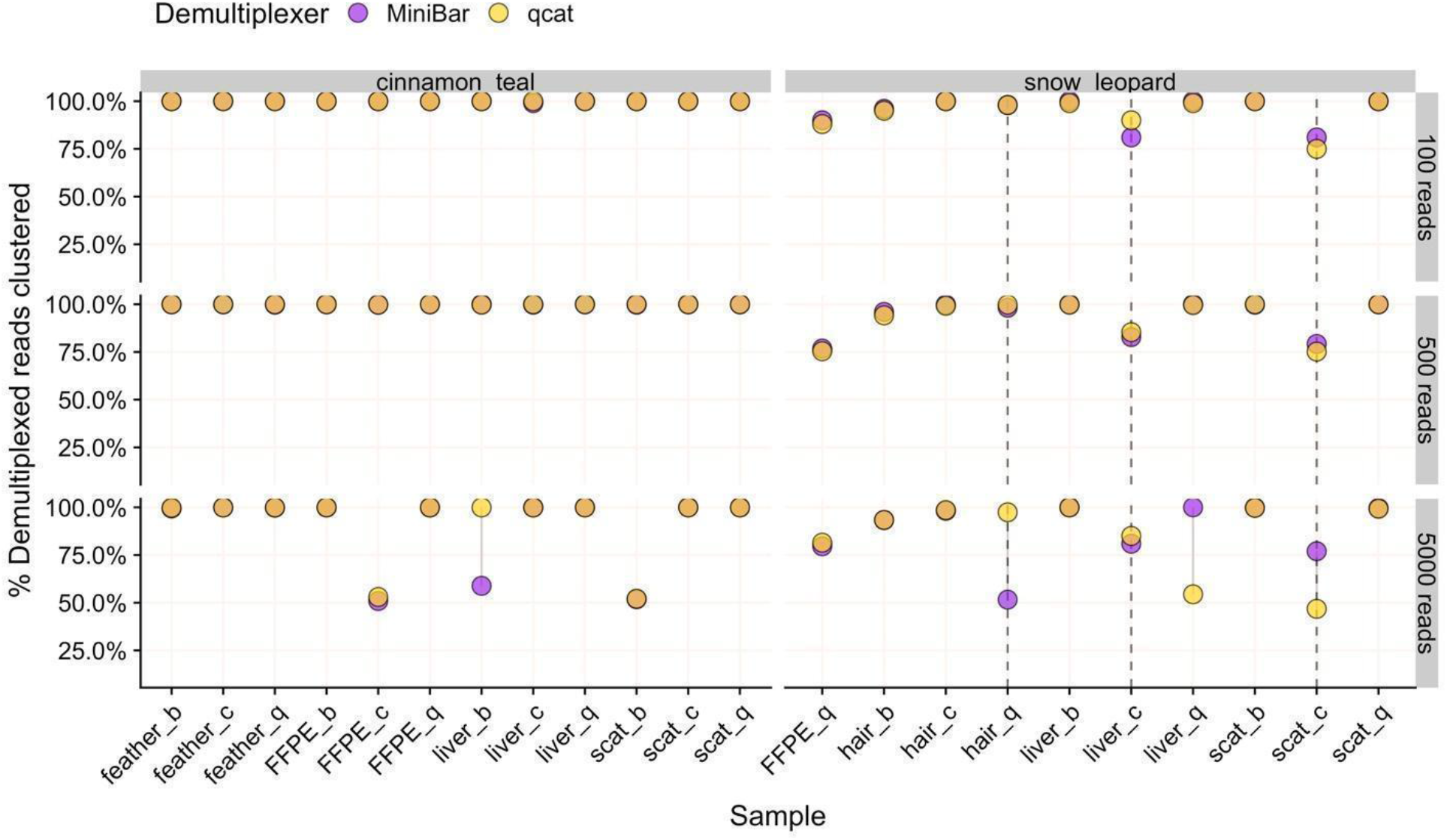
The percent of demultiplexed reads used to generate the final consensus sequence for 100R, 500R, and 5KR subsets for each species. Samples are labeled by tissue type and extraction method (b=biomeme, c=chelex, q=qiagen). Points are linked by a grey line to show difference in values from demultiplexers. Overlapping areas in orange indicate similar results for Minibar and qcat analyses. Vertical dashed lines indicate samples with cinnamon teal contamination.

In the remaining 18 data subsets, there were two categories: 1) samples where fewer than 60% of reads were used for final consensus generation due to sequence error and 2) samples with clusters containing contaminant reads (Table S5). In 5KR subsets for three cinnamon teal (FFPE/Chelex, liver/Biomeme, scat/Biomeme) and two snow leopard (hair/Qiagen, liver/Qiagen) samples, the second largest isONclust cluster contained reads that best match the same species as the majority cluster. While SPOA consensus sequences for these two clusters remained separate after cd-hit-est clustering, likely due to sequencing error (Table S5), species identification was successful for these five 5KR subset samples using only ~50% of the reads to build the consensus. In comparison, 100% of the reads clustered for the 100R and 500R subsets for these samples, suggesting that the 5KR subsample contained slightly more variation in read quality than the smaller subsets.

We detected low to medium levels of cinnamon teal reads in three snow leopard samples: hair/Qiagen, scat/Chelex, and liver/Chelex, where the full set of demultiplexed reads contained 3.9%, 22.0%, and 14.4% teal reads, respectively. There were no teal contaminant reads, and hence no teal read clusters, in the snow leopard hair/Qiagen sample for all subsets. In contrast, the proportions of reads used to generate final consensus for all subsets of the snow leopard scat/Chelex and liver/Chelex samples were reduced to 75-85% of reads (Table S5). Recovery of DNA Barcoding PCR (Round 1) products was low for these two samples. However, our pipeline’s filtering and clustering procedures were able to correctly identify these samples as snow leopard because reads with high sequence errors and contaminant reads were not included in downstream analysis. There were no cinnamon teal reads in the rest of the snow leopard samples, and no snow leopard reads in any cinnamon teal samples.

### Consensus sequence generation

The average proportion of reads used and consensus sequence lengths were comparable between sample types, extraction methods, subsets and demultiplexers (Table 1, Table S6). In general, SAIGA retained similar proportions of reads to generate consensus sequences across samples extracted by the Biomeme and Chelex methods as compared to the gold standard Qiagen-extracted samples (Fig. 3, Table 1, Table S6). In two cases, greater proportions of reads were used for the snow leopard liver and hair samples extracted with the Biomeme and Chelex protocols compared to the Qiagen-extract of the same tissue type. For samples where the consensus sequence length differed by demultiplexer, MiniBar subsets produced slightly longer sequences than qcat subsets (Fig. S6).

**Table 1:** Average and standard deviation (sd) for percent sequence similarity to Sanger sequence, length of matching nucleotides, and number and percent of demultiplexed reads used for the final consensus sequence from 100R, 500R, or 5KR read subsets demultiplexed with MiniBar or qcat. Statistics were calculated across all tissue types and extraction method samples.

| Subset | Demultiplexer | Average % ID (sd) | Average alignment length (bp) (sd) | Average number of clustered reads (sd) | Average % clustered reads (sd) |
| --- | --- | --- | --- | --- | --- |
| 100 reads per sample (100R) | MiniBar | 99.99 (0.05) | 421.05 (0.21) | 97.5 (5.8) | 97.50% (0.06) |
|  | qcat | 100 (0.00) | 420.5 (0.86) | 97.45 (6.01) | 97.45% (0.06) |
| 500 reads per sample (500R) | MiniBar | 99.97 (0.11) | 421.09 (0.43) | 484.5 (35.77) | 96.90% (0.07) |
|  | qcat | 100 (0.00) | 420.82 (0.59) | 483.68 (38.32) | 96.73% (0.08) |
| 5,000 reads per sample (5KR) | MiniBar | 99.88 (0.24) | 421.18 (0.8) | 4411.14 (916.69) | 88.22% (0.18) |
|  | qcat | 99.95 (0.18) | 420.41 (0.85) | 4456.14 (939.87) | 89.12% (0.19) |

### Validation of sample species identity

The average sequence similarity between MinION consensus sequences and their corresponding Sanger sequence was highly accurate (>99.29% match) and remarkably consistent across sample type, extraction method, subset, and demultiplexer (Fig. 4, Table 1). There was slightly more variation in sequence similarity across 5KR subsets, with the overall lowest percent sequence match (99.29%) obtained in these subsets for the cinnamon teal scat/Biomeme sample. This sample also had lower read cluster proportions (Fig. 3) and the greatest loss in data after filtering (Fig. S5).

**Figure 4:**
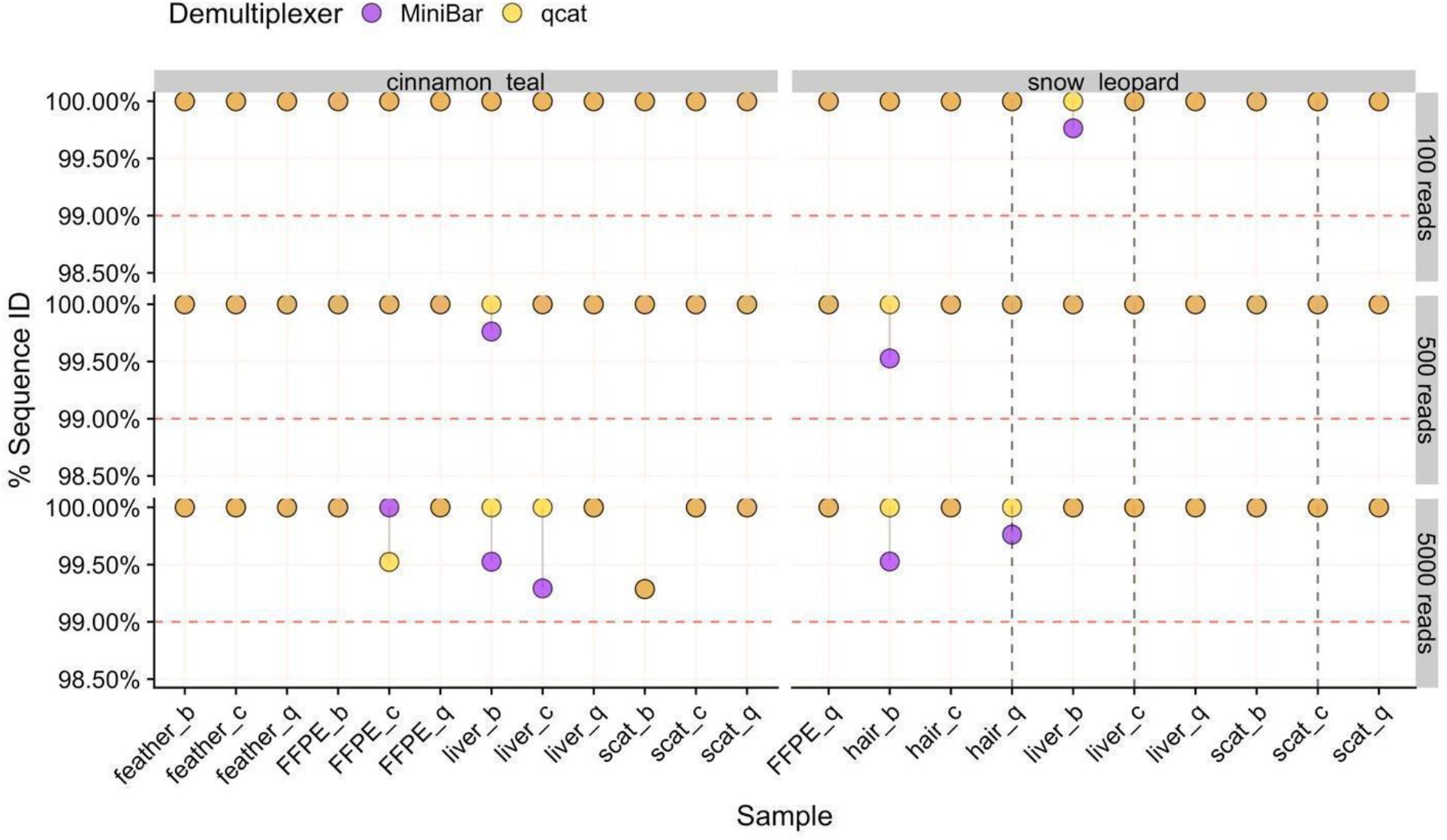
The percent sequence similarity of MinION consensus to Sanger sequence from Blast for 100R, 500R, and 5KR subsets for each species. Samples are labeled by tissue type and extraction method (b=biomeme, c=chelex, q=qiagen). Points are linked by a grey line to show difference in values from demultiplexers. Overlapping areas in orange indicate similar results for Minibar and qcat analyses. The horizontal dashed line is the 99% threshold for sequence similarity. Vertical dashed lines indicate samples with cinnamon teal read contamination.

The MinION consensus sequences from both MiniBar- and qcat-demultiplexed subsets extended into the Cytb primer region. We trimmed away the primers from both Sanger and MinION consensus sequences for Mafft alignment of all samples. The cinnamon teal alignment had 99.8% pairwise identity and 97.2% identical sites (n=84 sequences), while the snow leopard alignment had 99.9% pairwise identity and 98.6% identical sites (n=69 sequences). The MinION consensus and Sanger sequences for each animal mainly differed at the ends of the sequences and at homopolymeric regions of varying lengths within the sequence (Table S7, Fig. 5).

**Figure 5:**
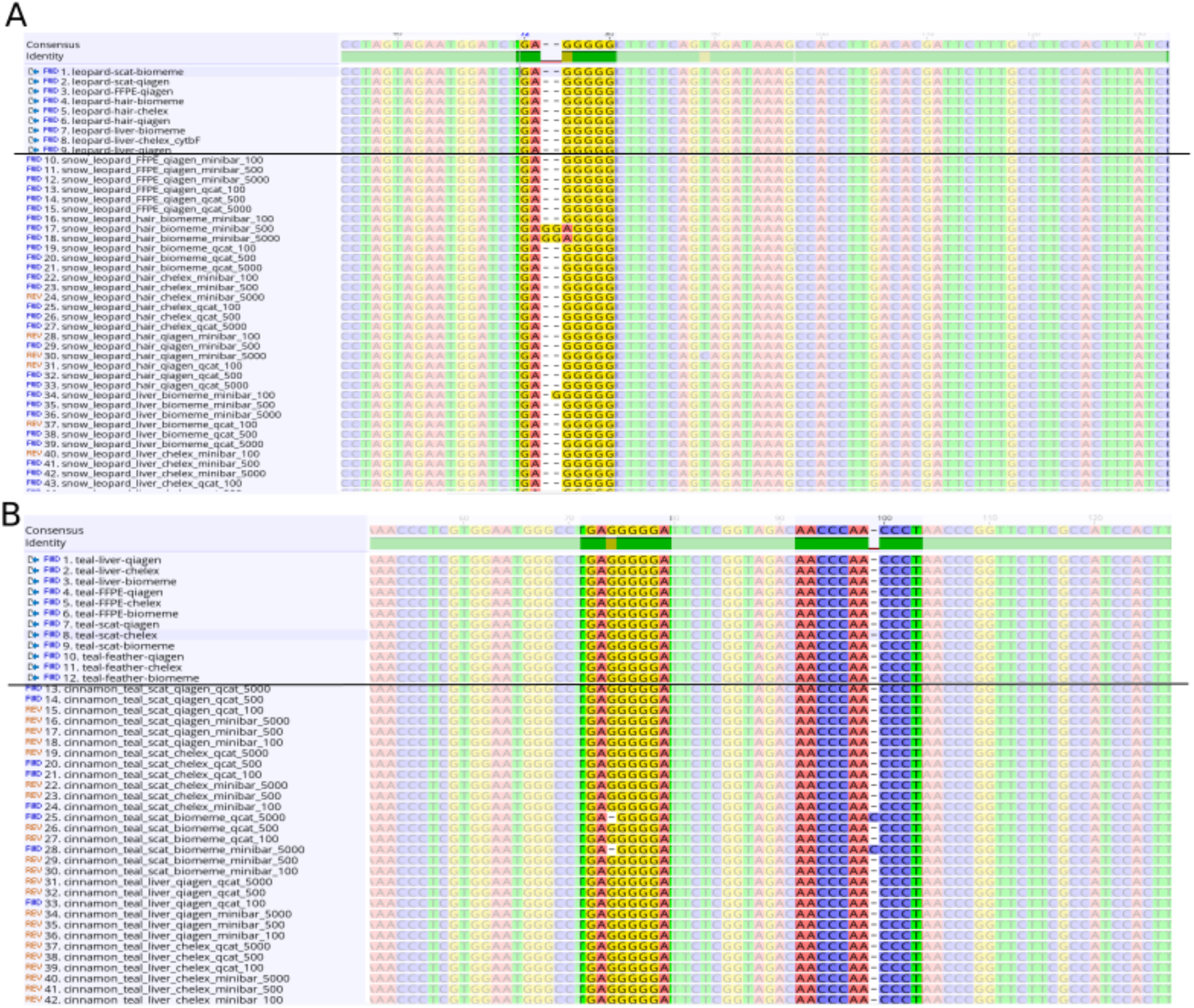
Screenshots of selected sections of the Mafft alignments for A) snow leopard and B) cinnamon teal showing nucleotide sites with differences between sequences in homopolymeric regions. Sanger sequences are listed above the black line and MinION consensus sequences below.

## Discussion

We demonstrate that a MinION-based DNA barcoding workflow can generate accurate consensus sequences from scat, hair, feather, and FFPE liver tissue samples, which are often considered challenging for molecular studies. The ability to use field-friendly DNA extraction protocols with these sample types will help to overcome logistical challenges, such as the need for cumbersome or expensive equipment, for molecular field research. The accuracy of our species identifications is on par with previous MinION DNA barcoding studies and pipelines (Pomerantz et al., 2018; Srivathsan et al., 2018, 2019; Krehenwinkel, Pomerantz, Henderson, et al., 2019; Maestri et al., 2019). For all tissue types, extraction methods, and subsets tested with our pipeline, we obtained high quality reads and a consensus sequence that matched >99.29% and at least 419/421 bp to the Sanger sequence for each sample. Although Oxford Nanopore’s goal is the “analysis of any living thing, by anyone, anywhere,” major barriers to its use are ease of sample processing, complicated data analysis, and cost. The results of our study help to reduce these barriers.

### Field-friendly protocols for wildlife samples expands conservation applications with the MinION

We show that the Chelex and Biomeme extraction methods can be used to generate highly accurate MinION consensus sequences, similar to Qiagen extraction methods, even with low starting DNA concentrations. Our PCR amplicon purification and library prep protocols resulted in libraries of sufficient purity; cellular debris or contaminants present in the Chelex and Biomeme extracts did not affect sequencing of the Cytb amplicons. Although the field-friendly DNA extracts had low DNA concentrations overall, amplification was successful for all samples, including scat (known for containing PCR inhibitors), hair and feather (low DNA quantities), and FFPE tissue, from which DNA is generally difficult to amplify.

Formalin can cause DNA fragmentation, cross-linking, subsequent sequence artifacts and altered base pairs (Do & Dobrovic, 2015; Einaga et al., 2017). As artifacts are randomly distributed, they should not affect the final Sanger sequence if sufficient starting template is used (Srinivasan, Sedmak, & Jewell, 2002; Quach, Goodman, & Shibata, 2004). Indeed, we accurately sequenced Qiagen-extracted DNA from FFPE samples, and further show that amplifiable DNA was successfully isolated from FFPE tissue using Chelex and Biomeme extraction methods.

### SAIGA: A DNA barcoding bioinformatics pipeline for new MinION users

We developed the SAIGA bioinformatics pipeline with a read clustering and consensus calling approach using software that were specifically designed for long-read and error-prone sequence data (isONclust, SPOA, Medaka). SAIGA performed successfully and consistently with as few as 100 reads per sample, allowing researchers to reduce sequencing time and cost per sample (e.g., multiplexing more samples). Like other studies investigating read coverage requirements, species identification accuracy still met our requirements but dropped slightly for the larger subset (5KR) (Pomerantz et al., 2018; Krehenwinkel, Pomerantz, Henderson, et al., 2019). Further, SAIGA options allow users to explore parameters and provide informative data quality checks and statistics throughout the pipeline. All software components are freely available and the pipeline structure allows for integration of new software in the future.

Our results show that both qcat and MiniBar correctly demultiplex reads between samples in a sequence run and across multiple runs on a flow cell. Due to the very stringent demultiplexing parameters, the majority of raw data loss occurred during read assignment. More relaxed settings reduce raw read loss, but increase the chance of including incorrectly assigned reads or reads with higher sequencing error. Srivathsan et al. (2019) and Maestri et al. (2019) noted similar magnitudes of read loss with ~76% and ~53.6% of reads lost after demultiplexing, respectively; other MinION DNA barcoding publications have not reported this statistic. Despite the read loss, MiniBar- and qcat-demultiplexed reads performed well based on all our metrics for accurate species identification. Both demultiplexers tend to under-trim reads, which is preferred since potentially useful regions of the amplicon for distinguishing species are lost from over-trimmed reads. Although the consensus accuracy of qcat results was slightly higher than MiniBar results, we prefer Minibar for its flexibility to analyze non-ONT index sequences. Customized indexes are less expensive than ONT indexes and can be lyophilized for field use.

Measuring the proportion of clustered filtered reads used for consensus sequence generation provides a benchmark for detecting sequencing error and potential contamination. For example, SAIGA created separate SPOA consensus sequence clusters for some samples even though these clusters produce the same species identification result. Lowering the sequence similarity threshold in cd-hit could force the sequences to form a single cluster. However, for the purpose of validating SAIGA, we used very stringent sequence similarity thresholds to reduce species identification bias from sequence error. Using this measure, we also show that SAIGA can handle low to medium amounts of laboratory contamination (~4-20% reads of total subsample) from relatively distinct species in samples without affecting final species identification since contaminant reads were successfully filtered out during the clustering process. Since contaminant teal reads had the correct indexes used for the three snow leopard samples, contamination likely occurred during library preparation rather than from mis-assignment of reads during demultiplexing. These snow leopard samples were either difficult to amplify during the Barcoding PCR (scat/Chelex) or had low recovery of indexed PCR product used in the sequencing run (hair/Biomeme and liver/Chelex). The contamination risk for these samples was likely exacerbated by the two-step PCR protocol and low starting DNA concentration and/or purity. Further development is needed to adapt this workflow and pipeline for mixed species samples, for which it may be more difficult to differentiate between true sample species and laboratory contaminants.

### Cost-effective strategies for field implementation

Each field-friendly method has its advantages and disadvantages. The Chelex method is cheap and the resin can be transported at room temperature, but requires heating equipment and the Chelex solution must be kept cool (4°C) once prepared. The Biomeme kit is room temperature stable and self-contained. However, it is more expensive than both the Chelex resin and Qiagen kits ($15/sample versus $0.17 and $3, respectively) and yielded lower DNA concentrations compared to the Qiagen kit.

We show that qcat and MiniBar can correctly assign reads to samples within and between runs, which reduces costs by allowing multiple sequence runs per flow cell. Future experiments can also scale up by sequencing more samples per flow cell because relatively few reads per sample are required for a consistent, accurate consensus (e.g. Srivathsan et al., 2019). For the Cytb barcode amplified in this study, reads were sequenced at a rate of ~100,000 reads per ~10 minutes. Sufficient sequence data for species barcoding can therefore be obtained rapidly depending on the barcoding gene length and number of samples. We also reduced the volumes of the ONT PCR index per sample by 50% to lower costs and maximize the ONT kit.

## Conclusions

Portable sequencing technology and field-friendly protocols have incredible potential to overcome institutional and geographical obstacles that impede genetic analyses in wildlife conservation and animal health. The methods described here provide an easy-to-follow workflow using field-friendly DNA extraction methods that can be used for preserved and non-invasively collected wildlife sample types to produce high-quality consensus sequences for species identification. Future studies are necessary to develop additional field-friendly protocols to further reduce the need for cold chain requirements, scale up sample processing, and tackle samples of mixed species, which will help to increase the opportunities for implementation.

## Acknowledgements

Funding was provided by the G. Unger Vetlesen Foundation. We thank Nina Vasiljevic and Rob Ogden for sharing their library preparation protocol and valuable discussions for our informatics pipeline, Batya Nightingale for lab assistance, and two anonymous reviewers for helpful comments.

## Author Contributions

AS and MCWL contributed equally to the project. AS, MCWL, DM, SP, and TS designed the study and interpreted the data. SP and MCWL developed SAIGA. AS conducted the lab work. MCWL performed the bioinformatics analysis. All authors contributed to writing the draft and gave final approval for publication.

## Data Availability

A representative Sanger sequence for both species is available on GenBank (MN823069-70), and MinION fastq files (basecalled, demultiplexed, and filtered) are available on NCBI Short Read Archive (BioProject: PRJNA594927, accessions: SRR10678113-SRR10678156). Raw MinION sequence data is available on the EBI European Nucleotide Archive (ERP119594).

