## Supplemental Material for "MinION-based DNA barcoding of preserved and non-invasively collected wildlife samples"

### Supplementary Materials

**Table S1:** Tailed Cytb primers used for indexing PCR. Tail sequence shown in purple/underlined font.

| Primer name | Primer sequence |
|---|---|
| mcb 398T | TTTCTGTTGGTGCTGATATTGCTACCATGAGGACAAATATCATTCTG |
| mcb 869T | ACTTGCCTGTCGCTCTATCTTCCCTCCTAGTTTGTTAGGGATTGATCG |

**Table S2:** DNA extraction concentrations for samples extracted with Qiagen and Biomeme kits were measured with the Qubit™ dsDNA High sensitivity kit on the Qubit™ 4 Fluorometer.

| Species | Tissue type | Extraction Method | DNA (ng/µL) |
|---|---|---|---|
| Cinnamon teal | FFPE | Qiagen | 47.6 |
| | | Biomeme | 0.336 |
| Snow leopard | | Qiagen | 59.0 |
| | | Biomeme | 0.452 |
| Cinnamon teal | Feather | Qiagen | 0.8 |
| | | Biomeme | 0.151 |
| Snow leopard | Hair | Qiagen | 0.723 |
| | | Biomeme | 0.129 |
| Cinnamon teal | Fecal | Qiagen | 34.2 |
| | | Biomeme | 4.57 |
| Snow leopard | | Qiagen | 38.5 |
| | | Biomeme | 1.66 |
| Cinnamon teal | Liver | Qiagen | 49.3 |
| | | Biomeme | 13.9 |
| Snow leopard | | Qiagen | 17.0 |
| | | Biomeme | 0.07 |

**Table S3:** The total number of reads, average read length, and average read quality score (qs) for raw data per sequencing run, and the averages across all sequencing runs. For demultiplexed and filtered reads, the number of total reads and the proportion of reads remaining from the previous pipeline step are shown for reads demultiplexed with MiniBar and qcat.

| Run/ Flow cell | Raw reads | | | Demultiplexed reads | | | | Filtered demultiplexed reads | | | |
|---|---|---|---|---|---|---|---|---|---|---|---|
| | | | | MiniBar | | qcat | | MiniBar | | qcat | |
| | Total reads | Ave read length (bp) | Ave read qs | Total reads | % reads kept | Total reads | % reads kept | Total reads | % reads kept | Total reads | % reads kept |
| 1/FAL 19910 | 515,142 | 599.5 | 10.6 | 95,264 | 18.49 | 106,331 | 20.64 | 93,927 | 98.60 | 102,412 | 96.31 |
| 2/FAL 19910 | 1,015,433 | 600.6 | 10.4 | 250,261 | 24.65 | 146,538 | 14.43 | 231,793 | 92.62 | 134,768 | 91.97 |
| 1/FAL 19272 | 729,764 | 593.0 | 10.5 | 71,553 | 9.80 | 130,435 | 17.87 | 69,948 | 97.76 | 125,277 | 96.05 |
| 2/FAL 19272 | 751,086 | 592.7 | 10.3 | 76,145 | 10.14 | 85,876 | 11.43 | 73,535 | 96.57 | 81,513 | 94.92 |
| Ave | 752,856.3 | 596.5 | 10.5 | 123,305.8 | 15.77 | 117,295.0 | 16.09 | 117,300.75 | 96.39 | 110,992.5 | 94.81 |

**Table S4:** Experimental design for each tissue type, including flow cell, number of active pores in thee pre-run Mux scan, species, extraction method, ONT index, demultiplexing information, and Sanger sequences. For full MiniBar and qcat demultiplexed and filtered read datasets, we show NCBI Short Read Archive (SRA) accessions, average quality score (qs), mean read length, and number of reads per sample; averages are calculated at the bottom of the table. 100R, 500R, and 5KR subsets were generated from the full demultiplexed and filtered read files. See supplemental file Table_S4.xlsx.

**Table S5:** The proportion of top cd-hit cluster reads used for final consensus generation (versus remaining clusters) are shown for samples with 1) proportions that were lower than 60% due to sequence error or 2) contaminant reads. For this table, the reads in remaining isONclust clusters excluded clusters with fewer than 10% of filtered reads and were not used to build the consensus sequence for final species identification.

| Species | Tissue | Extraction | Subset | Demultiplexer | % reads used for final consensus | % reads in remaining isONclust clusters | Reason |
|---|---|---|---|---|---|---|---|
| Cinnamon teal | FFPE | Chelex | 5000 | MiniBar | 51% | 49% | Sequence error |
| | | | | qcat | 53% | 47% | |
| | Liver | Biomeme | 5000 | MiniBar | 59% | 41% | |
| | Scat | Biomeme | 5000 | MiniBar | 52% | 48% | |
| | | | | qcat | 52% | 48% | |
| Snow leopard | Hair | Qiagen | 5000 | MiniBar | 52% | 42% | |
| | Liver | Qiagen | 5000 | qcat | 54% | 45% | |
| | Liver | Chelex | 100 | MiniBar | 81% | 15% | Contamination (with cinnamon teal) |
| | | | 500 | MiniBar | 83% | 15% | |
| | | | | qcat | 85% | 15% | |
| | | | 5000 | MiniBar | 80% | 13% | |
| | | | | qcat | 85% | 12% | |
| | Scat | Chelex | 100 | MiniBar | 81% | 19% | |

| | | | | | | |
|---|---|---|---|---|---|---|
| | | | | qcat | 75% | 25% |
| | | | 500 | MiniBar | 79% | 21% |
| | | | | qcat | 75% | 25% |
| | | | 5000 | MiniBar | 77% | 23% |
| | | | | qcat | 77% | 23% |

**Table S6:** A comparison of the average and standard deviation (sd) for percent of demultiplexed reads used for the final consensus sequence from subsets of 100, 500, or 5,000 reads demultiplexed with MiniBar or qcat for each extraction method. Statistics were calculated across all tissue types.

| Subset | Demultiplexer | Extraction Method | Average % ID (sd) | Average alignment length (bp) (sd) | Average # of clustered reads (sd) | Average % clustered reads (sd) |
|---|---|---|---|---|---|---|
| 100 reads | MiniBar | Biomeme | 99.97 (0.09) | 421.14 (0.38) | 99.43 (1.51) | 99.43% (0.02) |
| | | Chelex | 100 (0) | 421 (0) | 94.43 (9.18) | 94.43% (0.09) |
| | | Qiagen | 100 (0) | 421 (0) | 98.5 (3.51) | 98.50% (0.04) |
| | qcat | Biomeme | 100 (0) | 420.71 (0.76) | 99.14 (1.86) | 99.14% (0.02) |
| | | Chelex | 100 (0) | 420.57 (0.79) | 95 (9.57) | 95.00% (0.1) |
| | | Qiagen | 100 (0) | 420.25 (1.04) | 98.13 (4.16) | 98.12% (0.04) |
| 500 reads | MiniBar | Biomeme | 99.9 (0.19) | 421.29 (0.76) | 496.86 (7.45) | 99.37% (0.01) |
| | | Chelex | 100 (0) | 421 (0) | 472.43 (46.36) | 94.48% (0.09) |
| | | Qiagen | 100 (0) | 421 (0) | 484.25 (40.63) | 96.85% (0.08) |

| | | | | | | |
|---|---|---|---|---|---|---|
| | qcat | Biomeme | 100 (0) | 421 (0) | 495.57 (10.85) | 99.11% (0.02) |
| | | Chelex | 100 (0) | 421 (0) | 471.14 (49.82) | 94.23% (0.1) |
| | | Qiagen | 100 (0) | 420.5 (0.93) | 484.25 (43.34) | 96.85% (0.09) |
| 5000 reads | MiniBar | Biomeme | 99.76 (0.31) | 421.14 (0.9) | 4309.43 (1063.71) | 86.19% (0.21) |
| | | Chelex | 99.9 (0.27) | 421.43 (1.13) | 4335.71 (928.23) | 86.71% (0.19) |
| | | Qiagen | 99.97 (0.08) | 421 (0) | 4566.13 (876.98) | 91.32% (0.18) |
| | qcat | Biomeme | 99.9 (0.27) | 420.43 (0.79) | 4606.57 (890.86) | 92.13% (0.18) |
| | | Chelex | 99.93 (0.18) | 420.29 (0.95) | 4166.43 (1175.02) | 83.33% (0.24) |
| | | Qiagen | 100 (0) | 420.5 (0.93) | 4578 (814.16) | 91.56% (0.16) |

**Table S7:** Sequence errors from Mafft alignments of MinION-generated consensus and Sanger sequences for each species.

| Species | Sequence error | Tissue type | Extraction Method | Demultiplexer | Subset |
|---|---|---|---|---|---|
| Cinnamon teal | Incorrect base (GA instead of AG) in a 4-G homopolymeric region in nucleotides 1 and 2 of the fragment (Fig. 5b) | Scat | Qiagen | qcat | 500R |
| | | Scat | Chelex | qcat | 100R |
| | | Scat | Biomeme | qcat | 5KR |
| | | Scat | Biomeme | MiniBar | 5KR |
| | | Liver | Qiagen | qcat | 100R |
| | | FFPE | Qiagen | qcat | 5KR |
| | | FFPE | Qiagen | qcat | 100R |
| | | FFPE | Chelex | qcat | 5KR |
| | | FFPE | Chelex | qcat | 100R |
| | | FFPE | Biomeme | qcat | 100R |
| | | Feather | Qiagen | qcat | 500R |
| | | Feather | Qiagen | qcat | 100R |
| | | Feather | Chelex | qcat | 5KR |
| | | Feather | Biomeme | qcat | 5KR |
| | Incorrect base (T → C) in 7-C homopolymeric region | Liver | Biomeme | MiniBar | 500R |
| | | Scat | Biomeme | qcat | 5KR |

| | | | | | |
|---|---|---|---|---|---|
| | Gap in 5-G homopolymeric region and C insertion in 3-C homopolymeric region (Fig. 5b) | Scat | Biomeme | qcat | 100R |
| | Gap in 5-G homopolymeric region | Liver | Biomeme | MiniBar | 5KR |
| | | FFPE | Chelex | qcat | 5KR |
| | Gap in 3-C homopolymeric region | Scat | Biomeme | qcat | 5KR |
| | | Scat | Biomeme | MiniBar | 5KR |
| | | Liver | Biomeme | MiniBar | 5KR |
| | | FFPE | Chelex | qcat | 5KR |
| | Repeat/insertion of TCA at the end of a 4-C homopolymeric region, that is followed by TCA | Liver | Chelex | MiniBar | 5KR |
| | Repeat/insertion of TCC that is followed by TCC | Liver | Biomeme | qcat | 5KR |
| Snow leopard | Insertions in a 5-G homopolymer region (Fig. 5a) | Hair | Biomeme | MiniBar | 500R |
| | | Hair | Biomeme | MiniBar | 5KR |
| | | Liver | Biomeme | MiniBar | 100R |
| | Incorrect base (T → C) in the middle of the fragment (Fig. 5a) | Hair | Qiagen | MiniBar | 5KR |
| | Incorrect base (T → C) at the end | Hair | Chelex | qcat | 5KR |
| | Incorrect base (T → C) at the end | Scat | Qiagen | qcat | 5KR |

**Figure S1:** Gel electrophoresis of Qiagen DNA extracts to assess DNA quality of all sample types for the two species: fresh frozen liver, scat, FFPE liver, and feather or hair. 1.25 µL of each Qiagen extract was run on a 1% gel in 1xTAE buffer with GelPilot 1 kb Plus Ladder (lanes M).

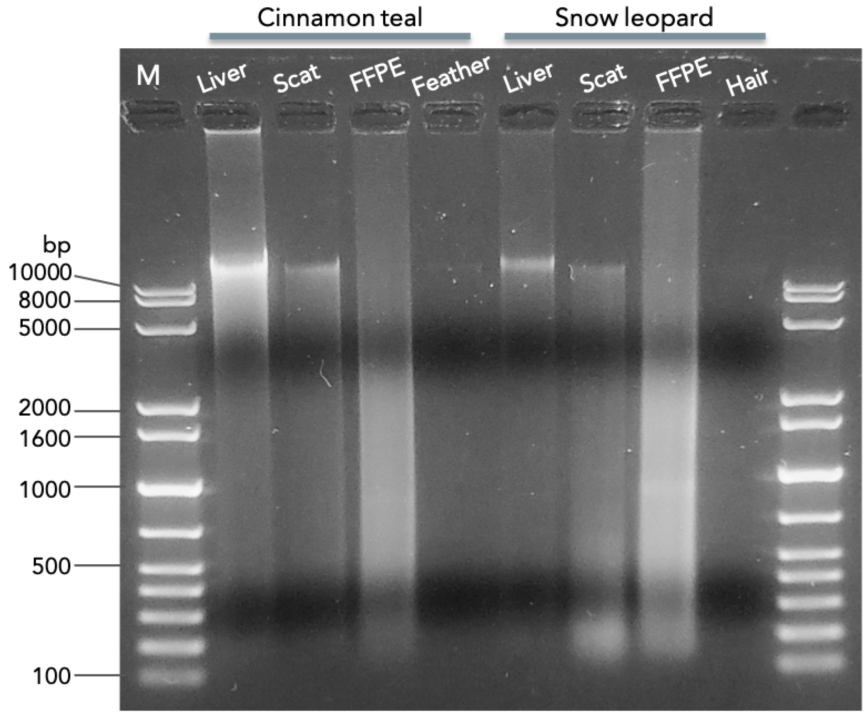

**Figure S2:** Gel electrophoresis of Qiagen, Chelex and Biomeme DNA extracts to assess DNA quality. Gel lanes are labeled with CT = Cinnamon teal, SL = Snow leopard, q = Qiagen, c = Chelex, and b = Biomeme. 10 μL of each Chelex extract, 1.25 μL of each Qiagen and one Biomeme extract were run on a 1% gel in 1xTAE buffer. Chelex samples had more low molecular weight nucleic acid. Chelex-extracted samples are difficult to assess because cellular debris that are not removed during extraction interfere with both quantification and quality measurements. Even after using 10ul of the undiluted Chelex extract for each sample (~80x more volume than was used in the DNA barcoding PCR), there was no obvious presence of high molecular weight nucleic acid for the Chelex extracts. It is possible that the visible low molecular weight fragments on the gel for Chelex samples includes RNA, however, SYBR Safe gel stain is not optimal for RNA binding. The single Biomeme extract had high enough concentration (>10ng/μL by Qubit) but was not visible on the gel, suggesting an overestimation of DNA concentration due to other contaminants in the extract or degradation since the sample was sequenced.

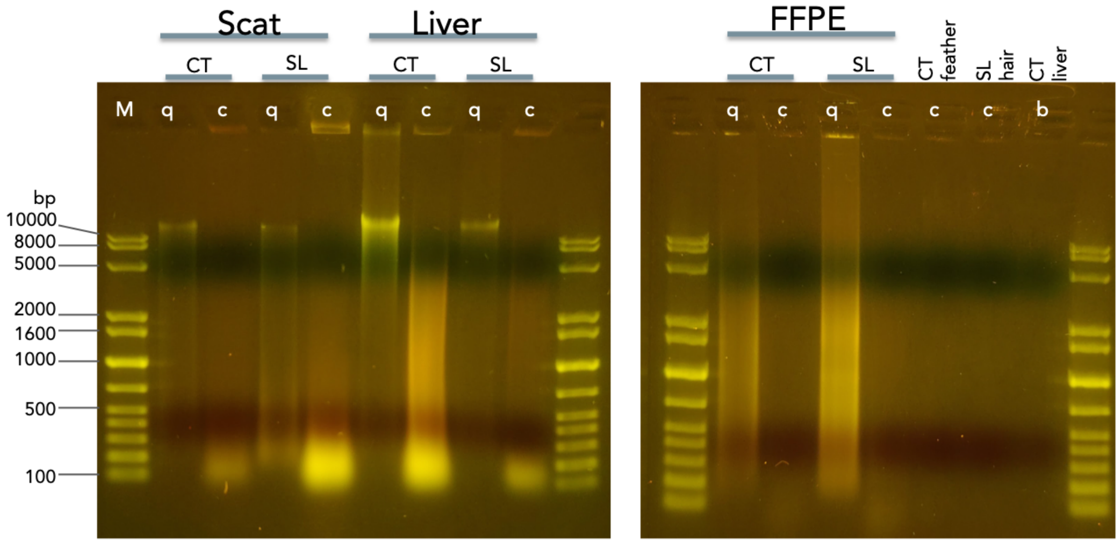

**Figure S3:** The number of reads generated per channel for each flow cell's sequencing run: A) 1st run on FAL19910, B) 2nd run on FAL19910, C) 1st run on FAL19272, and D) 2nd run on FAL19272. Plots generated by NanoPlot.

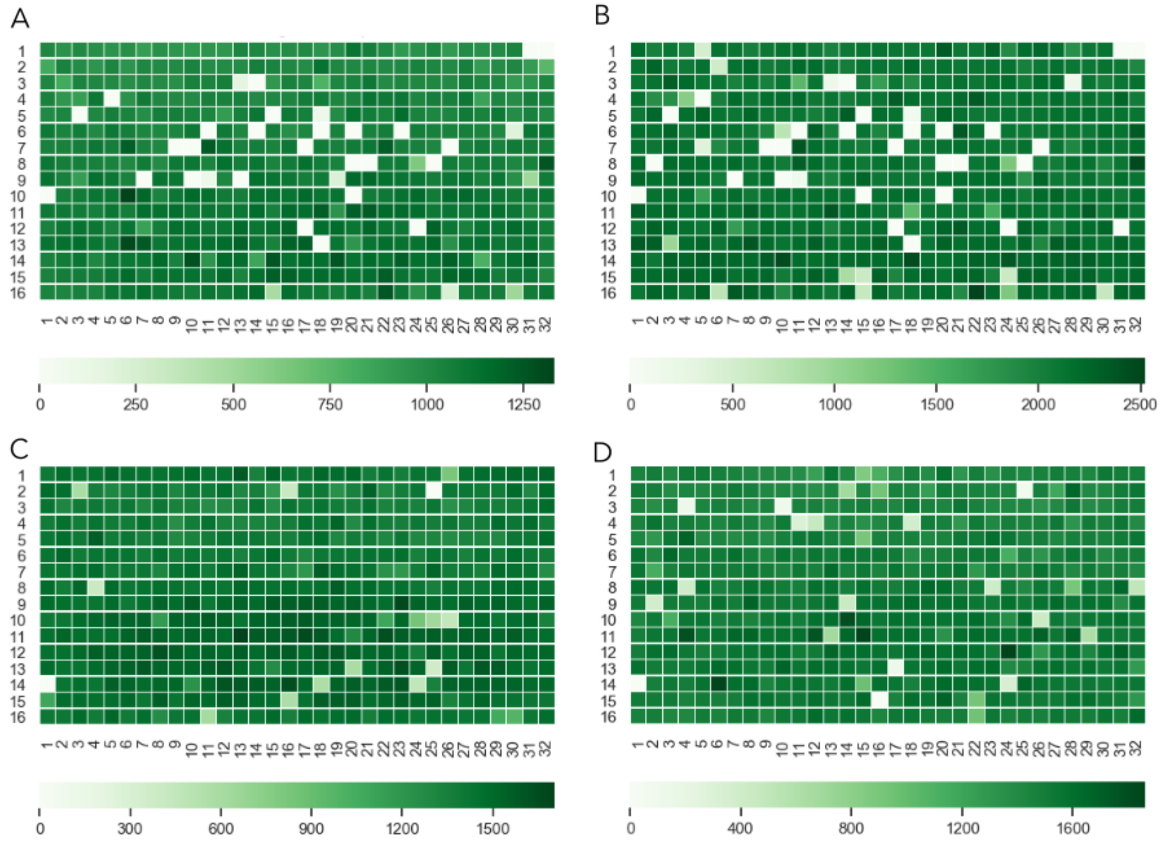

**Figure S4:** Violin plots of basecall quality for each flow cell's sequencing run duration (~1 hour). The widest region of the violin plot corresponds to basecall quality for the majority of the reads. Plots generated by NanoPlot.

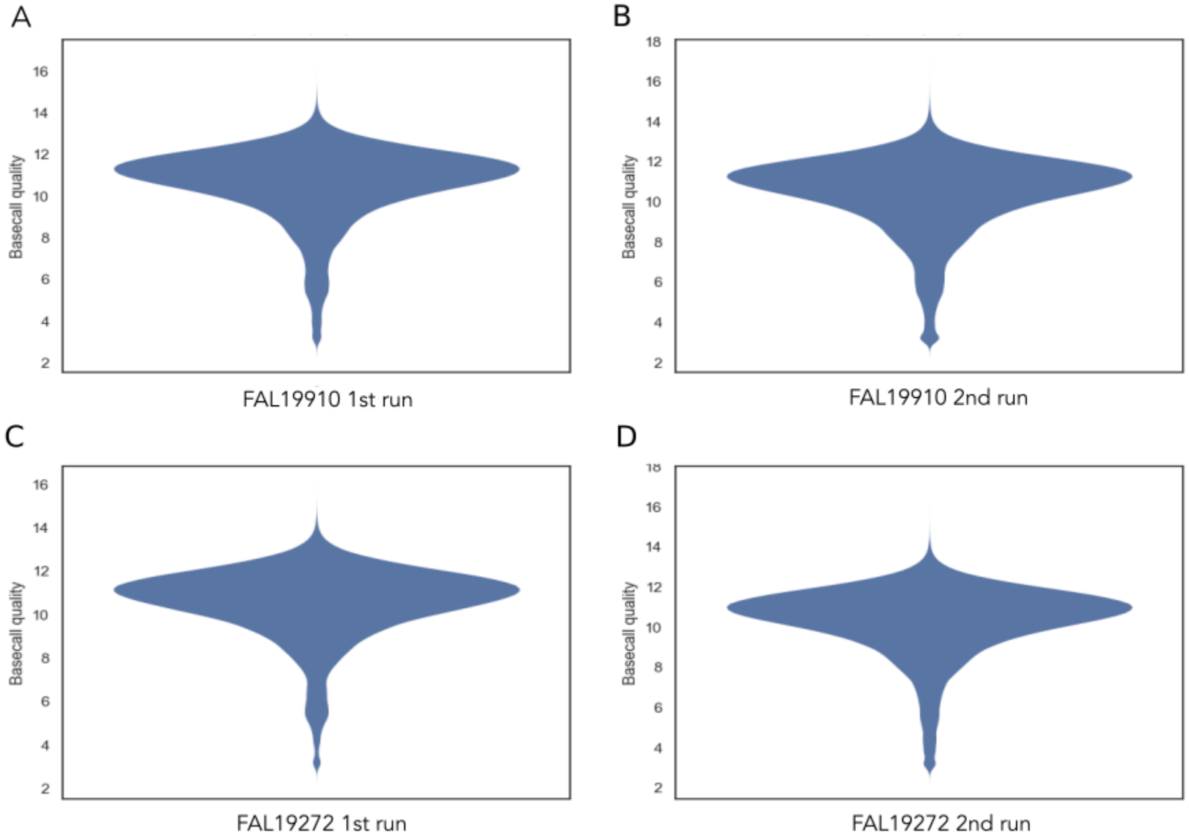

**Figure S5:** The proportion of demultiplexed reads remaining after filtering by read quality and length for each sample. Results are shown for MiniBar- and qcat-demultiplexed datasets. Samples are labeled by tissue type and extraction method (b=biomeme, c=chelex, q=qiagen). Points are linked by a black line to show difference in values from demultiplexers. Overlapping areas in orange indicate similar results for Minibar and qcat analyses.

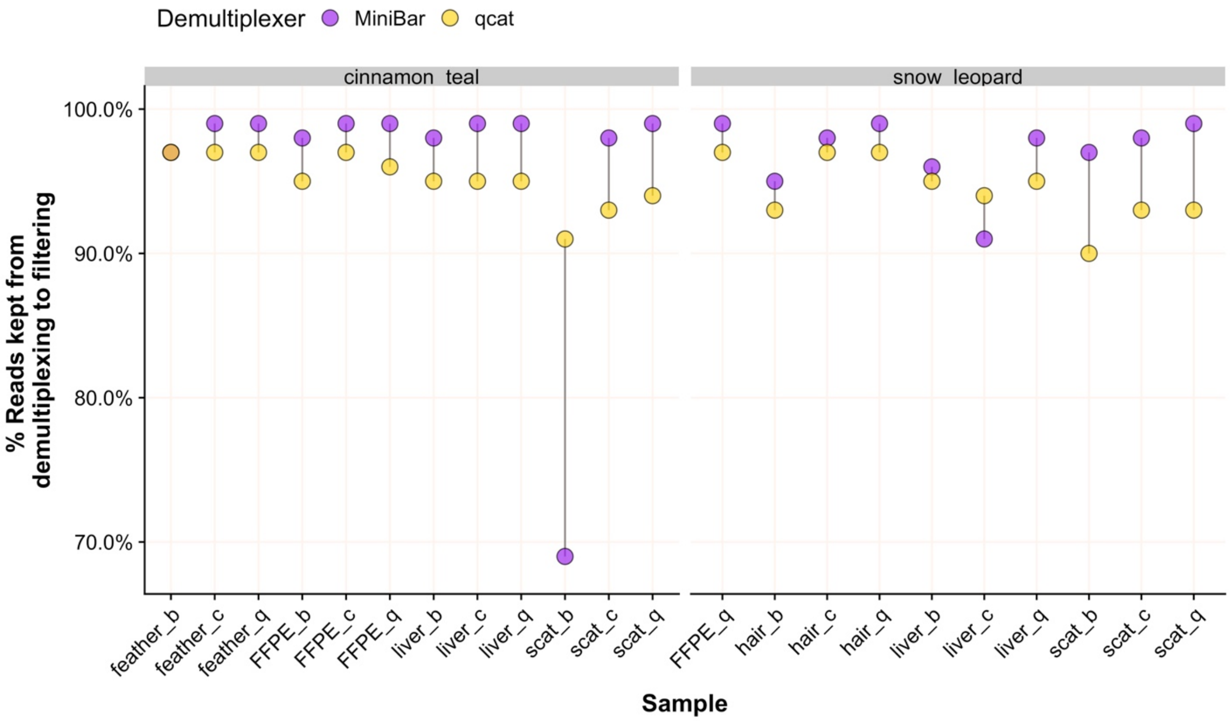

**Figure S6:** The number of nucleotides that match between MinION consensus to Sanger sequence from Blast for 100R, 500R, and 5KR subsets for each species. Samples are labeled by tissue type and extraction method (b=biomeme, c=chelex, q=qiagen). Points are linked by a grey line to show difference in values from demultiplexers. Overlapping areas in orange indicate similar results for Minibar and qcat analyses.

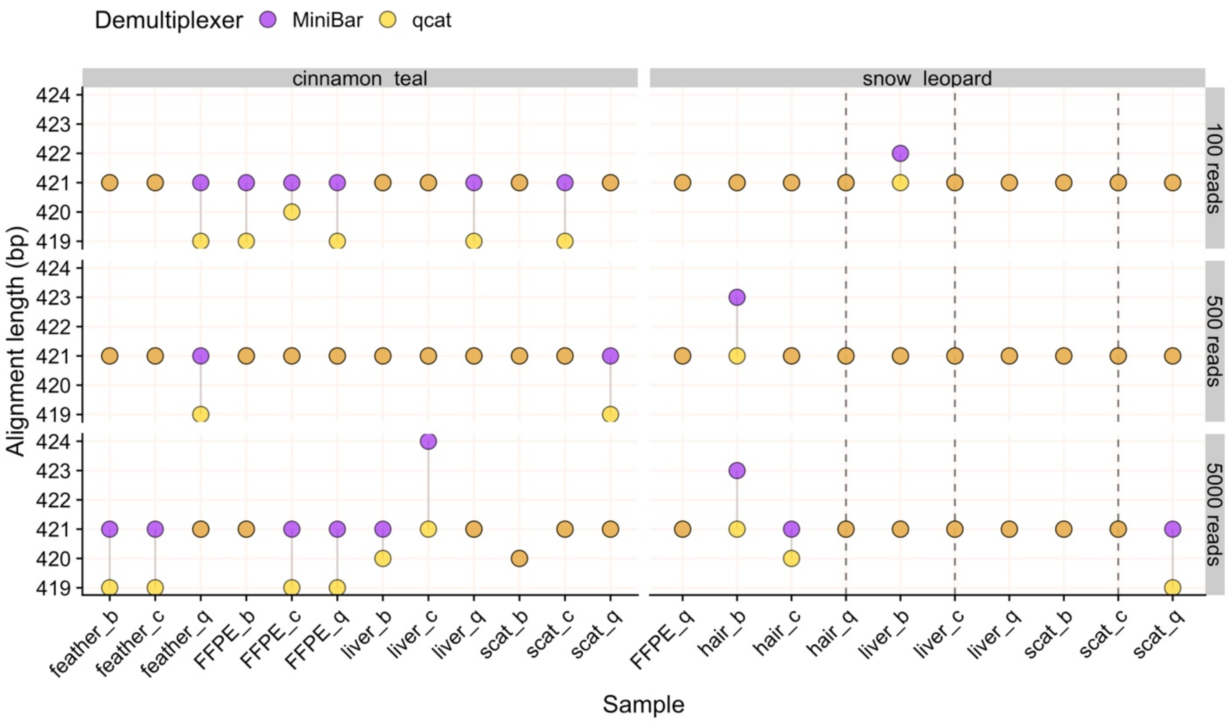
