## Appendix 1 for "MinION-based DNA barcoding of preserved and non-invasively collected wildlife samples"

### Appendix I

*Frozen liver samples*

1) DNA was extracted from a 3 $mm^3$cube of frozen liver using the QIAamp DNA Mini kit per manufacturer's instructions, and eluted in DNAse- and RNAse-free distilled water.

2) 190 µL of 5% Chelex reagent (dissolved in DNAse-free distilled water) and 10 µL of proteinase K (25mg/mL) were added to a 3 $mm^3$cube of frozen liver, followed by incubation at 55°C for 1 hour, and then 100°C for 20 minutes. Sample extracts were diluted 1:2 before being added to each PCR reaction for the DNA barcoding PCR (Round 1).

3) A 3 $mm^3$ cube of frozen liver was added to 0.5 mL of 3X Biomeme Lysis Buffer (BLB) and incubated at room temperature for 10 minutes. Next, the lysed sample was pumped through the Biomeme sample prep syringe (10 pumps). Subsequent washes were carried out with 0.5 mL Biomeme Protein Wash (1 pump), 1 mL Biomeme Wash Buffer (1 pump) and 1 mL Biomeme Drying Wash (1 pump), with a final drying step of 20 pumps of air. DNA was eluted off the column syringe in 0.5 mL Biomeme Elution Buffer (10 pumps).

*Formalin-fixed paraffin-embedded (FFPE) samples*

1) A total of 50 µm (10 x 5 µm thick sections) were processed following the Supplementary Protocol of the QIAamp® DNA FFPE Tissue kit (Qiagen, Germantown, MD, USA) using 320 µL of deparaffinization solution. Extraction was carried out according to the manufacturer's instructions. DNA was eluted in DNAse- and RNAse-free distilled water.

2) For the Chelex extraction, a total of 50 µm (10 x 5 µm thick sections) were processed using the Supplementary Protocol of the QIAamp DNA FFPE Tissue kit (Qiagen, Germantown, USA), with 320 µL of deparaffinization solution, and modified by replacing Buffer ATL with 190 µL of 20% Chelex and 10 µL proteinase K. The lower clear phase was kept as the DNA extract. For the DNA Barcoding PCR (Round 1), a 1:2 and 1:10 dilution of the snow leopard and cinnamon teal FFPE DNA extracts, respectively, were used in the PCR reactions.

3) For the Biomeme extraction, a total of 50 µm (10 x 5 µm thick sections) were processed following the Supplementary Protocol of the QIAamp® DNA FFPE Tissue kit (Qiagen, Germantown, MD, USA), with 320 µL deparaffinization solution, and modified to replace Buffer ATL with 500 µL BLB and 20 µL proteinase K (25 mg/mL). The lower clear phase was mixed in 0.5 mL of Biomeme Lysis Buffer (BLB) and incubated at room temperature for 10 minutes. The lysed sample was then pumped through the syringe (10 pumps), followed by 0.5 mL Biomeme Protein Wash (1 pump), 1 mL Biomeme Wash Buffer (1 pump), 1 mL Biomeme Drying Wash (1 pump), and a final drying step of 20 pumps of air. DNA was eluted through the syringe in 0.25 mL Biomeme Elution Buffer (10 pumps).

*Scat samples*

1) DNA was extracted from 0.2 g of frozen scat using the QIAamp® DNA Stool Mini Kit (Qiagen, Germantown, MD, USA) according to the manufacturer's instructions, and eluted in DNAse- and RNAse-free distilled water.

2) 190 µL of 20% Chelex reagent and 10 µL of proteinase K (25 mg/mL) were added to 0.2 g of frozen scat, followed by incubation at 55°C for 1 hour, and then 100°C for 20 minutes. For the DNA Barcoding PCR (Round 1), a 1:100 dilution of the snow leopard scat extract was used, and a 1:10 dilution of the cinnamon teal scat extract was used.

3) 1 mL of BLB was added to 0.2 g of frozen scat, hand shaken for 30 seconds and incubated for 30 minutes at room temperature. The lysed sample was then pumped through the syringe (10 pumps), followed by 0.5 mL Biomeme Protein Wash (1 pump), 1 mL Biomeme Wash Buffer (1 pump), 1 mL Biomeme Drying Wash (1 pump), and a final drying step of 20 pumps of air. DNA was eluted through the syringe in 0.25 mL Biomeme Elution Buffer (10 pumps).

*Hair and feather samples*
1) DNA was extracted from a small clump of snow leopard hair (10 x 3 x 3 mm) and the tips of 3 cinnamon teal feathers using the QIAamp® DNA Mini Kit (Qiagen, Germantown, MD, USA) according to the manufacturer's instructions, and eluted in DNAse- and RNAse-free distilled water.

2) 190 µL of 20% Chelex reagent and 10 µL of proteinase K (25 mg/mL) were added to a small clump of snow leopard hair (10 x 3 x 3 mm) and the tips of 3 cinnamon teal feathers, followed by incubation at 55°C for 1 hour, and then 100°C for 20 minutes. A 1:2 dilution of the extract was used for the DNA Barcoding PCR (Round 1).

3) 0.5 mL of BLB was added to a small clump of snow leopard hair (10 x 3 x 3 mm) and the tips of 3 cinnamon teal feathers, hand shaken for 30 seconds and incubated for 30 minutes at room temperature. The lysed sample was then pumped through the syringe (10 pumps), followed by 0.5 mL Biomeme Protein Wash (1 pump), 1 mL Biomeme Wash Buffer (1 pump), 1 mL Biomeme Drying Wash (1 pump), and a final drying step of 20 pumps of air. DNA was eluted through the syringe in 0.25 mL Biomeme Elution Buffer (10 pumps).
